# GlobalUsefulNativeTrees, a database of 14,014 tree species and their uses, supports synergies between biodiversity recovery and local livelihoods in landscape restoration

**DOI:** 10.1101/2022.11.25.517923

**Authors:** Roeland Kindt, Lars Graudal, Jens-Peter Lilleso, Fabio Pedercini, Paul Smith, Ramni Jamnadass

## Abstract

Tree planting has the potential to improve the livelihoods of millions of people as well as to support environmental services such as biodiversity preservation. Planting however needs to be executed wisely if benefits are to be achieved. We have developed the GlobalUsefulNativeTrees (GlobUNT) database to directly support the principles advocated by the ‘golden rules for reforestation’, including planting tree mixtures that maximize the benefits to local livelihoods and the diversity of native trees. Developed primarily by combining data from GlobalTreeSearch with the World Checklist of Useful Plant Species, GlobUNT includes 14,014 tree species that can be filtered for ten major use categories, across 242 countries and territories. In a subcontinental comparison GlobUNT revealed that Malesia had the highest useful tree species richness (3,349) and was also richest for materials (2,723), medicines (1,533), human food (958), fuel (734), environmental uses (632), social uses (614), animal food (443), poisons (322) and invertebrate food (266).

## Introduction

Trees play major functional roles in the world’s ecosystems where they protect biodiversity and are important carbon sequesters that mitigate climate change ^(1; 2)^. They also provide a wide range of socioeconomic benefits to billions of people ^(3)^ that include being important sources of nutrient-rich foods that support healthy diets ^(4; 5)^. By growing diverse food trees in their agroforestry systems, for example, smallholder farmers can achieve year-round nutritional security, while the sale of these foods supports broader healthful consumption ^(6)^. Such agroforestry systems are recognised as an important and relatively low-cost restoration mechanism to meet massive current forest landscape restoration targets (https://www.bonnchallenge.org) that can benefit hundreds of millions of people ^(7)^.

*Mosaic-type restoration* aims to restore or create a landscape of multiple land uses, including land uses that include trees ^(8)^. However, despite their enormous potential to generate positive impacts for livelihoods and environmental services, large-scale tree planting initiatives are prone to fail if not planned and executed wisely ^(9; 10)^. Tree species to be planted should be selected carefully, for example to avoid the biosafety risks associated with promoting invasive species ^(11; 12)^. At the same time, trees planted should contribute to local livelihoods and biodiversity – not just carbon sequestration, often the dominant consideration of the past ^(13)^.

A fundamental principle to delivering better the diverse tree portfolios essential for successful forest landscape restoration is to consider more specifically the uses of tree species for the local communities that are involved in trees’ planting and management. Significant win-win opportunities exist for driving forest landscape restoration adoption and improving local peoples’ livelihoods if proper consideration is given to the uses of the trees to be planted. This is because careful reference to trees’ uses that meets specific local needs is an important incentive for community involvement in restoration action ^(4; 14)^. What is needed especially is a knowledge of uses of native tree species whose planting and management in restoration activities is supportive of the twin goals of biodiversity conservation and livelihood improvement, avoiding some of the detrimental impacts of focusing on better-known exotic trees.

Concerns related to the failure of current tree-planting initiatives and the need to restore ecosystem functions and deliver diverse benefits to local communities and biodiversity have recently led to the formulation of the ‘10 golden rules for reforestation’ ^(15)^. These principles currently guide the formulation of a Global Biodiversity Standard (https://www.biodiversitystandard.org/), which aims to (a) assess impacts of tree planting programmes on biodiversity, and (b) provide mentoring and support to tree-planting practitioners for better livelihood and biodiversity outcomes. Among these principles, listed as Rule #6, is to ‘Select species to maximize biodiversity’ specifying that monocultures should be avoided wherever possible in the circumstances when planting is required to restore sites targeted for restoration. Furthermore, the same principle states that the planting mixtures should (a) maximize the number of native tree species, and (b) exclude invasive species.

Here, we describe the development of the GlobalUsefulNativeTrees (GlobUNT) database to support the application of useful native trees toward successful forest landscape restoration action. This new database is based primarily on combining information from two sources, the GlobalTreeSearch database ^(16)^, which documents the native country or territory of distribution of all known tree species globally, and the World Checklist of Useful Plant Species ^(17)^, which lists over 40,000 plants with their different documented human uses. The resulting GlobUNT database, which includes data for over 14,000 tree species, constitutes the largest available dataset on trees, their uses and their distributions.

Our new database allows users to select diverse assemblages of tree species native to chosen countries and territories that are useful for the provision of ranges of specific products and services. GlobUNT also has a range of extra functionalities that include the ability to generates summary tables of differences in tree species, genus and family richness at subcontinental and continental levels. This information is important for communicating the potential livelihood benefits provided by native tree species to policymakers and practitioners involved in the regional planning that is essential for effective forest landscape restoration action. To further explore the potential of GlobUNT for contributing to biodiversity conservation along with livelihood provision, we specifically analysed the subcontinental distributions of endemic and threatened useful tree species based on data on threat status taken from the Global Tree Assessment To further explore the potential of GlobUNT for contributing to biodiversity conservation along with livelihood provision, we specifically analysed the subcontinental distributions of endemic and threatened useful tree species based on data on threat status taken from the Global Tree Assessment ^(18; 19)^ (GTA; https://www.globaltreeassessment.org/).

## Results

### The global native distribution of useful tree species by major use categories

At the global level, the richness of useful tree species (*S*_*u*_) was 14,014, representing roughly one third (33.7%, Table 1) of all plant species with documented uses in the World Checklist of Useful Plant Species (WCUPS) and one quarter (24.2%, Table 2) of all tree species documented by GlobalTreeSearch. Only for one use category was the global *S*_*u*_ lower than 1000 (Invertebrate Food with 712); this was also the category in the WCUPS with lowest richness overall. Among the 14,014 species in the database, 64 species were listed for all ten use categories, from 56 genera and with five genera that included more than one species (*Cordia, Prosopis, Tarchonanthus, Vachellia* and *Ziziphus*). 118 species (0.8%) were listed for nine use categories and 209 (1.5%) for eight. 6,776 species (48.3%) only had one use category and 2,871 species (20.5%) had two.

**Table 1.** Species richness in GlobalUsefulNativeTrees for the twenty subcontinents and countries with highest richness overall.

| Level | Geography | Area | ALL | AF | EU | FU | GS | HF | IF | MA | ME | PO | SU |
| --- | --- | --- | --- | --- | --- | --- | --- | --- | --- | --- | --- | --- | --- |
| S | Malesia | 2,525 | 3,349 | 443 | 632 | 734 | 176 | 958 | 266 | 2,723 | 1,533 | 322 | 614 |
|  | Indo-China | 1,881 | 2,037 | 218 | 463 | 335 | 156 | 555 | 103 | 1,488 | 1,401 | 239 | 250 |
|  | Western South America | 3,721 | 1,883 | 145 | 280 | 131 | 121 | 320 | 47 | 1,033 | 1,303 | 95 | 108 |
|  | Indian Subcontinent | 4,118 | 1,823 | 246 | 443 | 314 | 162 | 502 | 115 | 1,285 | 1,470 | 223 | 242 |
|  | West-Central Tropical Africa | 4,041 | 1,668 | 360 | 592 | 571 | 476 | 819 | 154 | 1,257 | 1,283 | 314 | 354 |
|  | South Tropical Africa | 3,258 | 1,438 | 405 | 631 | 542 | 438 | 767 | 160 | 1,085 | 1,201 | 278 | 329 |
|  | Papuasias | 481 | 1,411 | 400 | 406 | 640 | 44 | 467 | 245 | 1,163 | 632 | 163 | 547 |
|  | Northern South America | 1,318 | 1,352 | 93 | 198 | 84 | 73 | 204 | 38 | 835 | 887 | 75 | 66 |
|  | East Tropical Africa | 1,655 | 1,258 | 380 | 587 | 552 | 405 | 687 | 163 | 928 | 1,019 | 256 | 293 |
|  | West Tropical Africa | 6,060 | 1,250 | 296 | 492 | 458 | 376 | 644 | 136 | 1,020 | 1,005 | 289 | 326 |
| C | Indonesia | 1,878 | 2,724 | 425 | 553 | 701 | 129 | 833 | 257 | 2,291 | 1,273 | 280 | 571 |
|  | Malaysia | 329 | 2,115 | 207 | 364 | 356 | 128 | 609 | 121 | 1,765 | 1,074 | 208 | 273 |
|  | Brazil | 8,358 | 1,772 | 107 | 202 | 96 | 100 | 249 | 31 | 1,059 | 1,143 | 70 | 74 |
|  | China | 9,425 | 1,594 | 146 | 551 | 217 | 215 | 381 | 78 | 975 | 1,105 | 156 | 141 |
|  | India | 2,973 | 1,591 | 232 | 409 | 293 | 143 | 466 | 106 | 1,147 | 1,290 | 211 | 229 |
|  | Thailand | 511 | 1,478 | 183 | 338 | 288 | 99 | 464 | 91 | 1,165 | 1,030 | 193 | 214 |
|  | Papua New Guinea | 453 | 1,361 | 395 | 398 | 634 | 40 | 452 | 242 | 1,136 | 603 | 160 | 540 |
|  | Colombia | 1,110 | 1,342 | 105 | 212 | 98 | 86 | 250 | 39 | 743 | 930 | 75 | 87 |
|  | Congo (DRC) | 2,267 | 1,228 | 284 | 477 | 466 | 367 | 632 | 123 | 942 | 978 | 255 | 277 |
|  | Myanmar | 653 | 1,226 | 162 | 322 | 232 | 100 | 369 | 71 | 926 | 932 | 170 | 177 |
| G | Global | - | 14,014 | 1,494 | 3,317 | 2,162 | 1,552 | 3,310 | 712 | 9,261 | 8,283 | 1,109 | 1,396 |
|  | WCUPS % | - | 34.8 | 33.7 | 36.9 | 85.5 | 29.8 | 47.0 | 68.4 | 67.8 | 31.1 | 36.8 | 53.8 |

**Table 2.** Patterns of endemism in GlobalUsefulNativeTrees for the twenty subcontinents and countries with highest richness overall.

| Level | Geography | ALL | ALL % | E1 | E1 % | NE1 | NE1 % | E2 | E2 % | NE2 | NE2 % |
| --- | --- | --- | --- | --- | --- | --- | --- | --- | --- | --- | --- |
| S | Malesia | 3,349 | 35.2 | 513 | 12.3 | 2,836 | 53.2 | 2,502 | 29.6 | 847 | 80.7 |
|  | Indo-China | 2,037 | 43.7 | 33 | 3.6 | 2,004 | 53.6 | 934 | 35.2 | 1,103 | 55.0 |
|  | Western South America | 1,883 | 19.4 | 96 | 3.4 | 1,787 | 25.9 | 1,456 | 16.2 | 427 | 59.6 |
|  | Indian Subcontinent | 1,823 | 57.6 | 316 | 29.9 | 1,507 | 71.6 | 876 | 45.8 | 947 | 75.7 |
|  | West-Central Tropical Africa | 1,668 | 48.9 | 41 | 7.0 | 1,627 | 57.6 | 1,547 | 47.1 | 121 | 99.2 |
|  | South Tropical Africa | 1,438 | 64.4 | 9 | 5.3 | 1,429 | 69.3 | 1,301 | 62.2 | 137 | 98.6 |
|  | Papuasias | 1,411 | 45.3 | 302 | 20.5 | 1,109 | 67.3 | 926 | 37.3 | 485 | 76.3 |
|  | Northern South America | 1,352 | 23.6 | 21 | 2.1 | 1,331 | 28.2 | 984 | 19.1 | 368 | 64.8 |
|  | East Tropical Africa | 1,258 | 58.6 | 54 | 14.9 | 1,204 | 67.5 | 1,091 | 55.4 | 167 | 94.9 |
|  | West Tropical Africa | 1,250 | 70.8 | 13 | 17.3 | 1,237 | 73.2 | 1,143 | 69.1 | 107 | 96.4 |
| C | Indonesia | 2,724 | 45.9 | 243 | 16.3 | 2,481 | 55.9 | 2,018 | 39.6 | 706 | 84.6 |
|  | Malaysia | 2,115 | 39.0 | 124 | 8.3 | 1,991 | 50.5 | 1,534 | 32.2 | 581 | 87.6 |
|  | Brazil | 1,772 | 20.2 | 383 | 9.3 | 1,389 | 29.7 | 1,541 | 18.2 | 231 | 66.8 |
|  | China | 1,594 | 34.5 | 269 | 12.6 | 1,325 | 53.2 | 393 | 16.2 | 1,201 | 54.7 |
|  | India | 1,591 | 60.7 | 165 | 25.3 | 1,426 | 72.3 | 694 | 47.8 | 897 | 76.5 |
|  | Thailand | 1,478 | 57.1 | 8 | 3.7 | 1,470 | 62.0 | 736 | 47.1 | 742 | 72.4 |
|  | Papua New Guinea | 1,361 | 47.6 | 287 | 22.4 | 1,074 | 67.8 | 906 | 39.7 | 455 | 78.4 |
|  | Colombia | 1,342 | 22.6 | 27 | 2.4 | 1,315 | 27.2 | 945 | 17.9 | 397 | 59.2 |
|  | Congo (DRC) | 1,228 | 60.0 | 13 | 7.8 | 1,215 | 64.6 | 1,135 | 58.1 | 93 | 100.0 |
|  | Myanmar | 1,226 | 60.5 | 12 | 6.6 | 1,214 | 65.8 | 464 | 50.3 | 762 | 69.1 |
| G | Global | 14,014 | 24.2 | 4,143 | 12.6 | 9,871 | 39.4 | 11,306 | 21.4 | 2,708 | 52.3 |

At a continental level, tropical Asia had the highest *S*_*u*_ overall with 5177 species (Supplementary Table 1), followed by Africa (3413), Southern America (3158) and temperate Asia (2118). Two continents had *S*_*u*_ below 1000, the Pacific (530) and Europe (299).

At a sub-continental level, *S*_*u*_ varied from 3349 to 4 (Supplementary Table 1). The value was highest overall in Malesia (comprised of Brunei Darussalam, Christmas Island, Cocos Islands, Indonesia, Malaysia, Philippines, Singapore and Timor-Leste) and lowest in Subarctic America (Greenland). *S*_*u*_ was also largest in Malesia for Materials (MA, 2723), Medicines (ME, 1533), Human Food (HF, 958), Fuel (FU, 734), Environmental Uses (EU, 632), Social Uses (SU, 614), Animal Food (AF, 443), Poisons (PO, 322) and Invertebrate Food (IF, 266) (Fig. 1). Indo-China (Cambodia, Lao People’s Democratic Republic, Thailand and Viet Nam) ranked second highest for MA (1488). West-Central Tropical Africa (Burundi, Cameroon, Central African Republic, Congo, The Democratic Republic of the Congo [DRC], Equatorial Guinea, Gabon, Rwanda and Sao Tomé and Principe) ranked highest for *S*_*u*_ in Gene Sources (GS, 476; this category of reported uses includes wild relatives of major crops which may be valuable for breeding programs) and second highest for HF (819) and PO (314). South Tropical Africa (Angola, Malawi, Mozambique, Zambia and Zimbabwe) ranked second highest in *S*_*u*_ for EU (631), GS (438) and AF (405). Papuasia (Papua New Guinea and the Solomon Islands) had the second highest *S*_*u*_ for FU (640), SU (547) and IF (245). The Indian Subcontinent (Bangladesh, Bhutan, India, Maldives, Nepal, Pakistan and Sri Lanka) ranked second for ME (1470). Western South America (the Plurinational State of Bolivia, Colombia, Ecuador and Peru) that ranked third overall in *S*_*u*_ (1883) never ranked second or third for separate use categories.

**Fig. 1:**
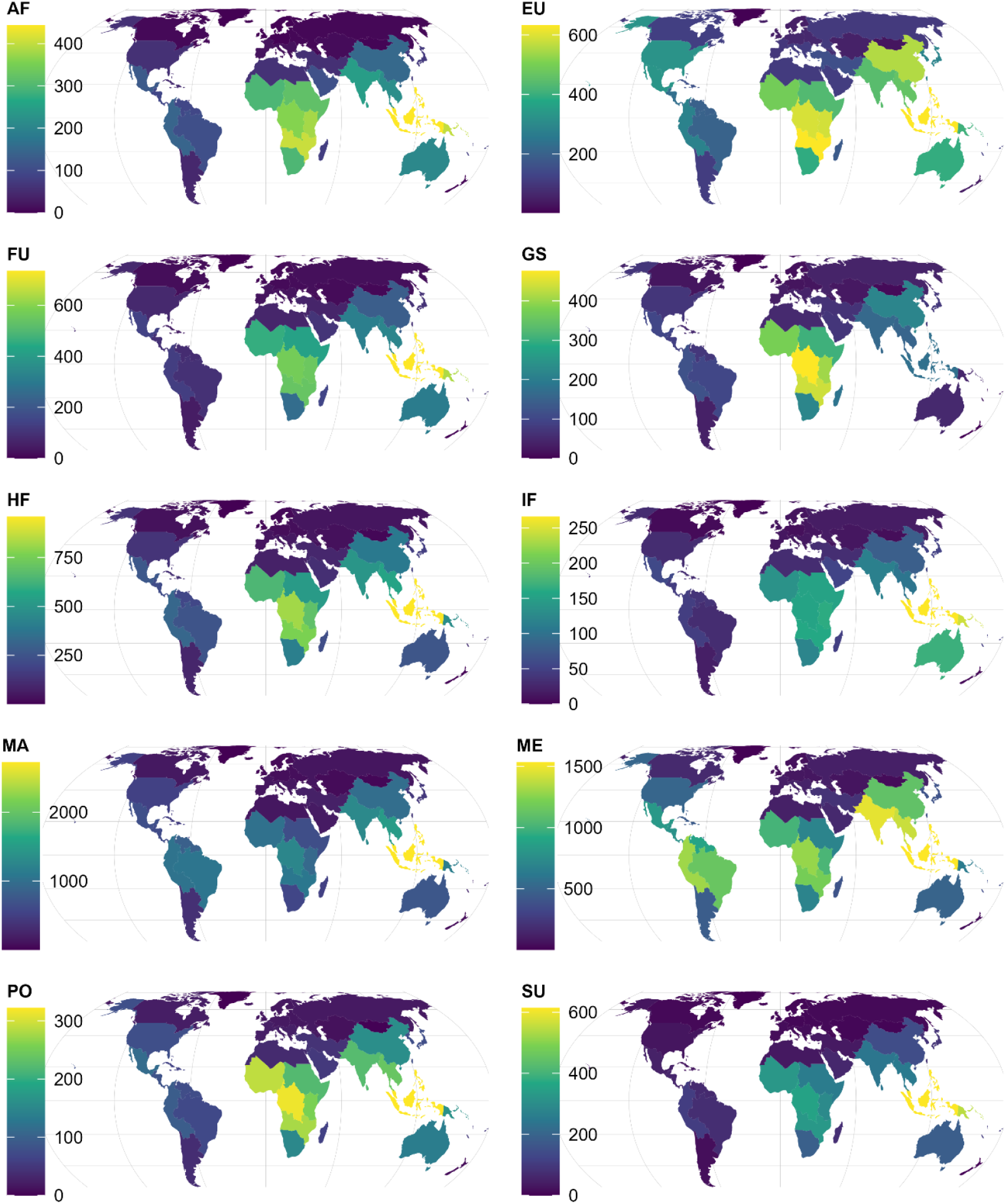
Subcontinental species richness for GlobalUsefulNativeTrees across different use categories. **AF**, Animal Food. **EU**, Environmental Uses. **FU**, Fuel. **GS**, Gene Sources. **HF**, Human Food. **IF**, Invertebrate Food. **MA**, Materials. **ME**, Medicines. **PO**, Poisons. **SU**, Social Uses (See (17) for definitions of reported uses). Supplementary Table 1 includes data for 242 countries and territories, 42 subcontinents and 8 continents for the 10 use categories. Map created in R with Equal Earth projection.

At a country/territory level, the species richness of native trees documented to be useful in GlobUNT (*S*_*u*_) ranged between 2724 and 1 (Supplementary Table 1). The Malesian countries of Indonesia and Malaysia ranked first and second in overall *S*_*u*_ with 2724 and 2115 species respectively. Indonesia also ranked first for MA (2291), HF (833), FU (701), SU (571), EU (553), AF (425), PO (280) and IF (257). Brazil, the most species-rich country in the GlobalTreeSearch database with 8791 species (Supplementary Table 1) ranked third overall in *S*_*u*_ (1772) and only had the same ranking for ME (1143) and otherwise ranked significantly lower for individual use categories with a highest sixth ranking for MA (1059).

Colombia that ranked second in GlobalTreeSearch (5943 species) only ranked eighth for GlobUNT (1342). India ranked first overall in *S*_*u*_ for ME (1290). The DRC ranked first overall in *S*_*u*_ for GS (367) and second highest for HF (632). China ranked second overall in *S*_*u*_ for EU (551, only two species lower than the best-ranked Indonesia) and GS (215).

Besides the countries and territories listed in Table 1, the United Republic of Tanzania had relatively high (ranking second or third) *S*_*u*_ for AF (318), EU (505), FU (470) and GS (344). Levels were also relatively high in Cameroon for GS (330) and in Nigeria for PO (253) and SU (283). Five countries not listed in Table 1 also had total *S*_*u*_ above 1000, Cameroon (1155), Mexico (1118), Peru (1106), the Plurinational State of Bolivia (1058) and the Philippines (1041).

At a plant family level, the total number in GlobUNT was 234 (GlobalTreeSearch includes 261 families). The twenty plant families with the highest *S*_*u*_ were the Fabaceae (1469), Rubiaceae (770), Myrtaceae (663), Malvaceae (565), Euphorbiaceae (541), Arecaceae (481), Lauraceae (460), Moraceae (419), Anacardiaceae (302), Sapotaceae (301), Annonaceae (294), Sapindaceae (289), Rutaceae (280), Phyllanthaceae (267), Apocynaceae (258), Rosaceae (251), Meliaceae (238), Dipterocarpaceae (226), Salicaceae (224) and Fagaceae (217). The total number of plant genera was 2599. The twenty genera with the highest *S*_*u*_ were *Ficus* (287), *Syzygium* (189), *Diospyros* (184), *Eucalyptus* (155), *Quercus* (117), *Terminalia* (99), *Acacia* (98), *Elaeocarpus* (96), *Garcinia* (96), *Croton* (94), *Prunus* (93), *Coffea* (90), *Pinus, Salix* (82), *Macaranga* (75), *Dombeya* (74), *Shorea* (74), *Commiphora* (73), *Magnolia* (69) and *Ilex* (67).

### The global native distribution of endemism and threat status of useful tree species shows x and y

To further explore the potential of GlobUNT for contributing to biodiversity conservation along with livelihood provision, we analysed the distributions of endemic and threatened trees.

When classifying endemism as a feature of useful tree species that is defined by them being native to a single country or territory only, the richness of endemic species (*S*_*e1*_) was largest in the subcontinents of Australia (557, Fig. 2, Supplementary Table 2) and the Western Indian Ocean (British Indian Ocean Territory, Comoros, Madagascar, Mauritius, Mayotte, Réunion and Seychelles, 515), followed by Malesia (513, Table 2), Brazil (383) and the Indian Subcontinent (316). The two countries with highest *S*_*e1*_ were Australia (557) and Madagascar (432), followed by Brazil (383) and Papua New Guinea (287). There were 64 countries where *S*_*u*_ included none of the single-country endemics (the country with the highest number of endemics in GlobalTreeSearch among these was Haiti with 163 species).

**Fig. 2:**
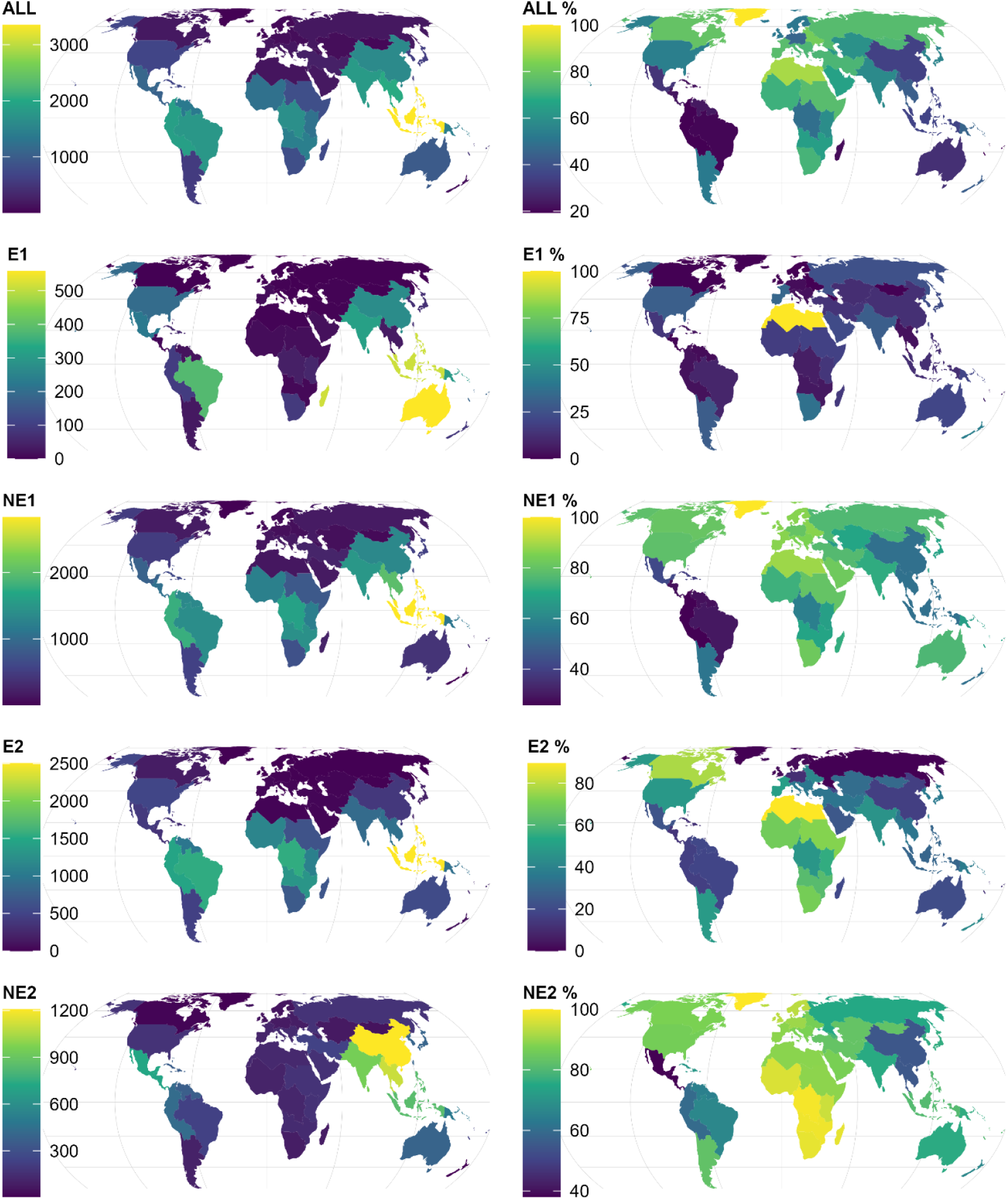
Subcontinental patterns of endemism for GlobalUsefulNativeTrees. **ALL**, All species in GlobUNT. **E1**, Species in GlobUNT native to one country only. **NE1**, Species in GlobUNT native to two countries or more. **E2**, Species in GlobUNT native to one continent only. **NE2**, Species in GlobUNT native to two continent or more. **ALL %-NE %**, Richness of left-hand panel expressed as percentages from the total number of species in GlobalTreeSearch in the same category. Supplementary Table 2 includes data on endemism for 242 countries and territories, 42 subcontinents and 8 continents. Map created in R with Equal Earth projection.

When defining endemic species by those useful tree species that were native to one continent only, the species richness of endemic species (*S*_*e2*_) was largest in Malesia (2502), West-Central Tropical Africa (1547) and Brazil (1541). Among the continents, the percentage of continent-endemics in GlobUNT versus those in GlobalTreeSearch was highest in Africa (34.5%), Northern America (28.7%) and tropical Asia (27.6%). The lowest percentages were in the Pacific (10.4%), Southern America (12.2%) and Europe (16.8%) (Supplementary Table 2).

Checking species that are not endemic to the continent showed that the largest numbers were in China (1202), India (897) and Viet Nam (858) (Table 2, Fig. 2).

The most widely distributed useful tree species across subcontinents were *Dodonaea viscosa* (28 subcontinents, 7 continents), *Ximenia americana* (24, 7), *Sophora tomentosa* (23, 7), *Hibiscus tiliaceus* (22, 7), *Pisonia aculeata* (22, 7), *Tephrosia purpurea* (20, 5), *Thespesia populnea* (20, 7), *Suriana maritima* (19, 7), *Avicennia marina* (18, 5), *Trema orientale* (18, 5) and *Vitex trifolia* (18, 6).

Defining threatened species as those with the IUCN Red List categories of Critically Endangered (CR), Endangered (EN) or Vulnerable (VU) resulted in the subcontinent of Malesia hosting the highest number of threatened species (*S*_*THR*_ = 382; Table 3; Fig. 3) with the Western Indian Ocean ranked second (245). These subcontinents showed the same ranking for CR (60 and 55 species, respectively) and EN (111 and 108, respectively). For VU, Malesia ranked first once again (211 species), but the Indian subcontinent ranked second (111). Among the subcontinents listed in Table 3, percentages of useful threatened tree species were highest in the Indian subcontinent for the categories of CR (27.3%) and EN (32.6%).

**Table 3.**
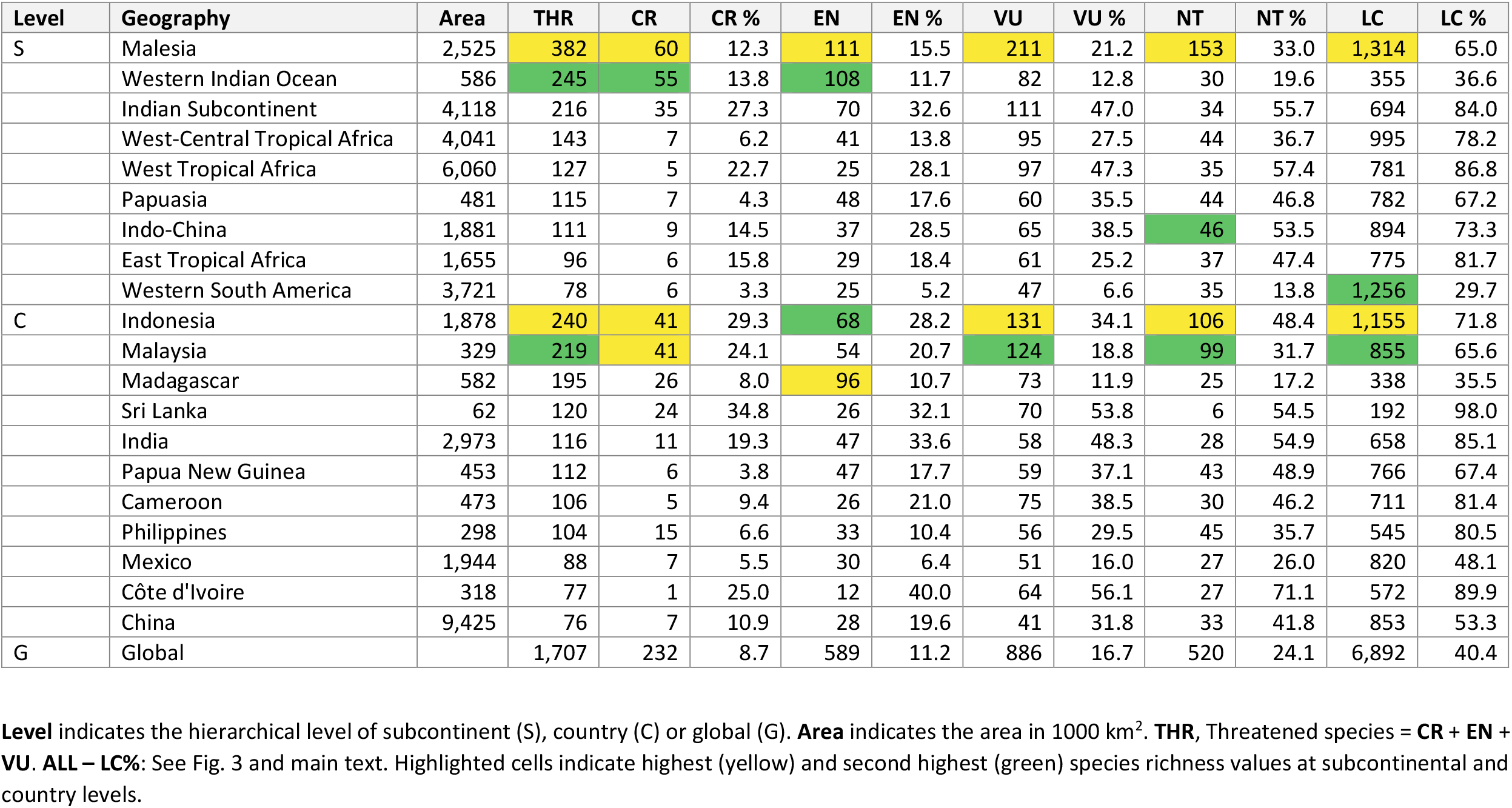
Patterns of threat status for species in GlobalUsefulNativeTrees for the twenty subcontinents and countries with highest numbers of threatened species.

| Level | Geography | Area | THR | CR | CR % | EN | EN % | VU | VU % | NT | NT % | LC | LC % |
| --- | --- | --- | --- | --- | --- | --- | --- | --- | --- | --- | --- | --- | --- |
| S | Malesia | 2,525 | 382 | 60 | 12.3 | 111 | 15.5 | 211 | 21.2 | 153 | 33.0 | 1,314 | 65.0 |
|  | Western Indian Ocean | 586 | 245 | 55 | 13.8 | 108 | 11.7 | 82 | 12.8 | 30 | 19.6 | 355 | 36.6 |
|  | Indian Subcontinent | 4,118 | 216 | 35 | 27.3 | 70 | 32.6 | 111 | 47.0 | 34 | 55.7 | 694 | 84.0 |
|  | West-Central Tropical Africa | 4,041 | 143 | 7 | 6.2 | 41 | 13.8 | 95 | 27.5 | 44 | 36.7 | 995 | 78.2 |
|  | West Tropical Africa | 6,060 | 127 | 5 | 22.7 | 25 | 28.1 | 97 | 47.3 | 35 | 57.4 | 781 | 86.8 |
|  | Papuasias | 481 | 115 | 7 | 4.3 | 48 | 17.6 | 60 | 35.5 | 44 | 46.8 | 782 | 67.2 |
|  | Indo-China | 1,881 | 111 | 9 | 14.5 | 37 | 28.5 | 65 | 38.5 | 46 | 53.5 | 894 | 73.3 |
|  | East Tropical Africa | 1,655 | 96 | 6 | 15.8 | 29 | 18.4 | 61 | 25.2 | 37 | 47.4 | 775 | 81.7 |
|  | Western South America | 3,721 | 78 | 6 | 3.3 | 25 | 5.2 | 47 | 6.6 | 35 | 13.8 | 1,256 | 29.7 |
| C | Indonesia | 1,878 | 240 | 41 | 29.3 | 68 | 28.2 | 131 | 34.1 | 106 | 48.4 | 1,155 | 71.8 |
|  | Malaysia | 329 | 219 | 41 | 24.1 | 54 | 20.7 | 124 | 18.8 | 99 | 31.7 | 855 | 65.6 |
|  | Madagascar | 582 | 195 | 26 | 8.0 | 96 | 10.7 | 73 | 11.9 | 25 | 17.2 | 338 | 35.5 |
|  | Sri Lanka | 62 | 120 | 24 | 34.8 | 26 | 32.1 | 70 | 53.8 | 6 | 54.5 | 192 | 98.0 |
|  | India | 2,973 | 116 | 11 | 19.3 | 47 | 33.6 | 58 | 48.3 | 28 | 54.9 | 658 | 85.1 |
|  | Papua New Guinea | 453 | 112 | 6 | 3.8 | 47 | 17.7 | 59 | 37.1 | 43 | 48.9 | 766 | 67.4 |
|  | Cameroon | 473 | 106 | 5 | 9.4 | 26 | 21.0 | 75 | 38.5 | 30 | 46.2 | 711 | 81.4 |
|  | Philippines | 298 | 104 | 15 | 6.6 | 33 | 10.4 | 56 | 29.5 | 45 | 35.7 | 545 | 80.5 |
|  | Mexico | 1,944 | 88 | 7 | 5.5 | 30 | 6.4 | 51 | 16.0 | 27 | 26.0 | 820 | 48.1 |
|  | Côte d'Ivoire | 318 | 77 | 1 | 25.0 | 12 | 40.0 | 64 | 56.1 | 27 | 71.1 | 572 | 89.9 |
|  | China | 9,425 | 76 | 7 | 10.9 | 28 | 19.6 | 41 | 31.8 | 33 | 41.8 | 853 | 53.3 |
| G | Global |  | 1,707 | 232 | 8.7 | 589 | 11.2 | 886 | 16.7 | 520 | 24.1 | 6,892 | 40.4 |

**Fig. 3:**
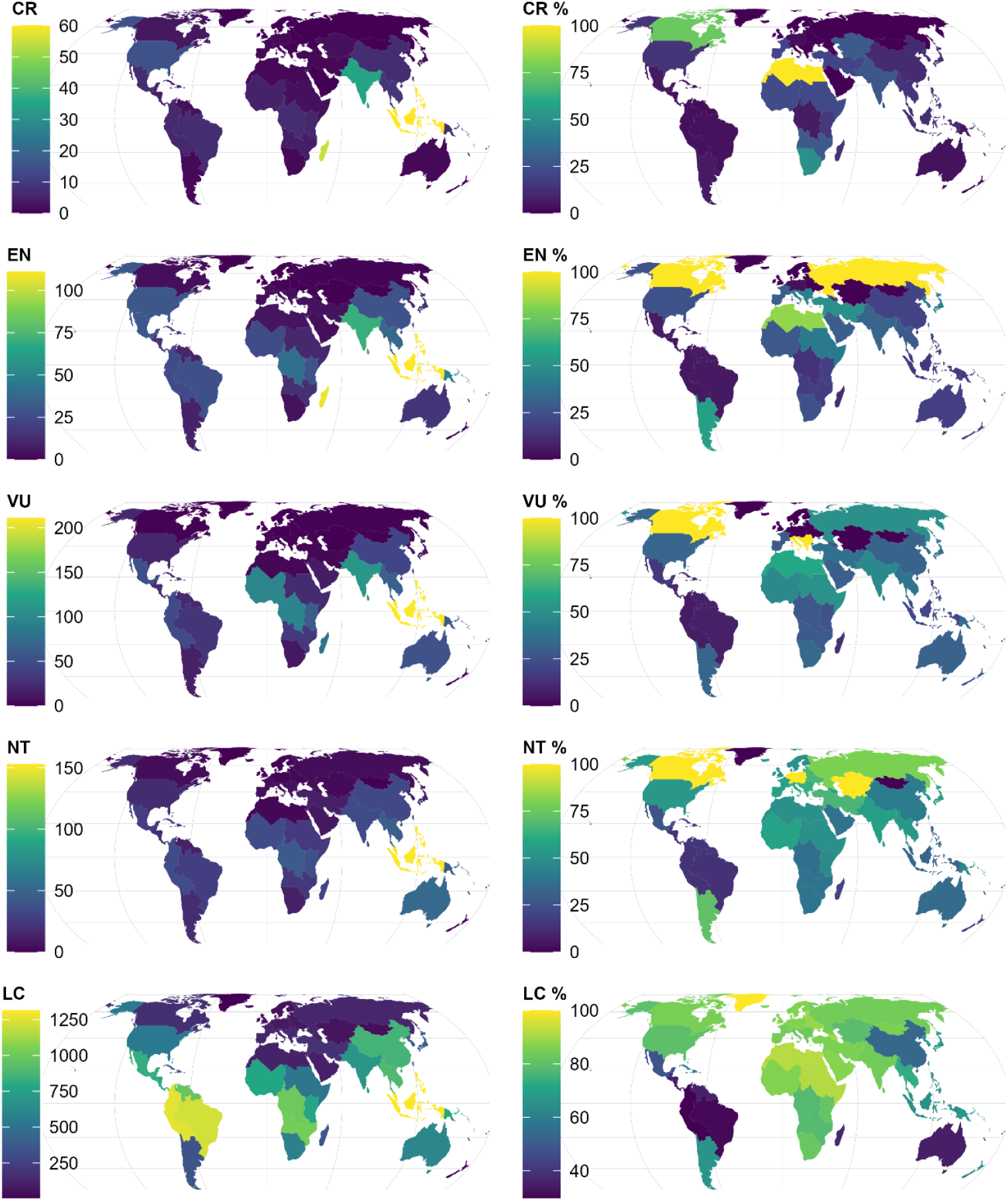
Subcontinental patterns of threats for GlobalUsefulNativeTrees. **CR**, Critically Endangered species. **EN**, Endangered species. **VU**, Vulnerable species. **NT**, Near Threatened species. **LC**, Species of Least Concern. **CR %-LC %**, Richness of left-hand panel expressed as percentages from the total number of species in GlobalTreeSearch in the same category. Supplementary Table 3 includes data on threats for 242 countries and territories, 42 subcontinents and 8 continents. Map created in R with Equal Earth projection.

The Malesian countries of Indonesia and Malaysia contained the highest numbers of threatened species (240 and 219 species, respectively). Madagascar had the highest number for EN (96). A country that was not mentioned earlier, Sri Lanka, ranked fourth overall in numbers of threatened species and also had more than twice the number of CR species than the fifth ranked India.

For species of Least Concern (LC), percentages of GlobUNT/GlobalTreeSearch were above fifty for most subcontinents and countries listed in Table 3. Exceptions were Western South America (29.7%), Madagascar (35.5%), the Western Indian Ocean (36.6%) and Mexico (48.1%).

Globally, percentages of threatened species were below 17% for the different categories for threatened species, whereas the percentage for LC was 40.4%.

## Discussion

Current tree-planting initiatives for forest landscape restoration are failing because they do not sufficiently consider the needs of local communities that they rely on for planting and tending the trees, such that the trees that are planted are not well cared for and do not survive to maturity. The range of species selected for use in restoration is furthermore limited, with an emphasis on exotic species that do not support biodiversity. To help address these concerns, the ‘10 golden rules for reforestation’ were recently developed ^(15)^. Among these ‘golden rules’ there is an emphasis on maximising native tree biodiversity and addressing local community needs to support success.

Our compilation of data primarily from GlobalTreeSearch and WCUPS (see methods on selected species and sources of information on tree uses) indicates that almost a quarter of all known trees have been assigned specific uses and that nearly a third of all useful plants are trees. These statistics highlight both the potential and limitations of focusing on useful tree species within global tree conservation schemes, thus highlighting the continued need for *in situ* conservation for tree species without known uses but also showing the scope for conservation-through-use of tree species with demonstrable value to humans.

Even if we focus solely on country endemics (12.6% of species represented in GlobUNT) or threatened species (12.9% listed by GlobUNT), the biodiversity conservation potential is not insignificant with over 4000 and 1700 species respectively representing win:win opportunities for human use and biodiversity conservation. Therefore, from a global perspective, it is possible to select planting mixtures from GlobUNT that satisfy the criteria of the ‘golden rules’ of favouring native species and also including endemic and threatened species (GlobUNT further includes hyperlinks to the GlobalTree Portal from where information is available on published IUCN Red List data and those of regional or national assessments as documented in the ThreatSearch database). At the same time, the selected assemblages will provide useful products and services. Furthermore, the database can allow for prioritizing based on desired services. However, a trade-off does exist between the number of species grown in landscape mosaics and their viable population sizes, and therefore tree densities should not pass thresholds that prevent their long term survival or enable connectivity ^(20)^.

GlobUNT is not the only database that allows selections of useful trees (overviews are provided elsewhere ^(21; 22)^ and country-specific manuals and tools also exist (see for example the RELMA-ICRAF Useful Trees series available via http://apps.worldagroforestry.org/usefultrees/index.php). However, when comparing the database with a selection of 27 databases with species-specific details for tree species that are available from the Agroforestry Species Switchboard ^(23)^, GlobUNT was the database that included the highest number of useful tree species (Supplementary Table 4). When considering all species listed in GlobalTreeSearch, only the obvious WCUPS, the Useful Tropical Plants Database ^(24)^ (57.8%) and wood density data available from the BIOMASS package ^(25)^ (51.0%) included more than half of the species included by GlobUNT. In addition to these databases, only the World Economic Plants in GRIN ^(26)^ (27.1%) and the Plant Resources of South East Asia ^(27)^ (24.6%) had close to or more than a quarter of species listed.

GlobUNT lists over 1000 species for potential use in 18 countries and over 300 species in 78 countries. For countries that have made pledges to bring degraded and deforested landscapes into restoration in response to the Bonn Challenge (https://www.bonnchallenge.org/pledges), GlobUNT provides 100 species or more for every country from Africa, tropical Asia, Northern America and Southern America except for Chile (Supplementary Table 5; combined, these 52 countries have pledged over 200 million ha).

Where GlobUNT lists 100 species or more, during the planning of tree planting projects it will be necessary to select subsets of species (conceptually such selection could be thought of as a local ecological filter selecting species from a regional species pool ^(28; 29)^). Filtering species can be done directly within GlobUNT by selecting the desired use categories. As described similarly in the ‘golden rules’ guidelines, country-specific checklists of native species can be used by local specialists to select a subset of species that is most suitable for a particular project site, for example by considering results from previous planting experiments ^(30)^. Since GlobUNT was developed in parallel with the previously mentioned *Switchboard*, specialists can readily access data from a large number of databases since this database provides verified hyperlinks to taxon-specific information across 45 information sources.

Malesia, Indonesia and Malaysia frequently obtained top rankings in the results. Brazil, the subcontinent and country of highest species richness in GlobalTreeSearch and the country of second-highest area coverage in Table 1, only got the third highest ranking overall and otherwise ranked lower except for the number of medicinal species and country-endemic species. The country did not feature among the countries with highest rankings for useful and threatened tree species (Table 3). We can only speculate why this is the case, but possibly the inclusion of more regional sources from the continents of Africa and tropical Asia within WCUPS could have biased species composition in WCUPS and subsequently in GlobUNT away from Brazil and Southern America.

In large countries that include several ecoregions (e.g., according to the *Ecoregions 2017* ^(31)^), it would be important as well to select species that match species assemblages of the natural vegetation; knowledge of native floras (many available online as for example via the World Flora Online website ^(32)^ or information from vegetation atlases (e.g, http://www.vegetationmap4africa.org) can be of help here. Similarly, where seed zonation maps have been developed (e.g., ^(33)^), or habitat distribution maps are available for individual tree species (e.g., ^(34; 35; 36; 37)^) these would be directly relevant in selecting species that match the environmental conditions of planting sites (and ideally future climatic conditions).

We want to underscore that our vision about usage of GlobUNT is not that the database is primarily used from a remote office as the principal method of deciding which species should be planted. On the contrary, tree planting projects wanting to avoid failure should, in addition to the ‘golden rules’ ^(15)^, also consider people-centred factors ^(14; 38)^. Previous recommendations for participatory selection remain highly valid, as recently repeated by ^(39; 40; 41)^. However, for any country where a project aims to implement tree planting schemes that aspire to maximise native tree biodiversity while addressing local community needs, GlobUNT will be a user-friendly source for practical information.

## Methods

### Species selection and distribution

The GlobalUsefulNativeTrees (GlobUNT) database includes 14,014 species. The identities of 13,947 species (99.5%) were obtained by matching species listed by GlobalTreeSearch ^(16; 42)^ (accessed on 8^th^ May 2022 for individual countries) with those listed in the World Checklist of Useful Plant Species ^(17)^ (WCUPS) by protocols documented below. Also included in GlobUNT were 62 species that were included in the WCUPS but not in the GlobalTreeSearch, but that otherwise had been included among 830 tree species prioritized for planting in the tropics and subtropics ^(43)^ (Top-830), that were listed in the Agroforestree database ^(44)^ (AFD) or listed within a selection of tree, bamboo and rattan species that are most widely planted in the tropics and subtropics ^(45)^. Details about these species are provided in Supplementary Table 6.

Four more species were added (*Acacia cincinnata, A. pachycarpa, Shorea javanica* and *Toona ciliata*) from GlobalTreeSearch and the Top-830 or AFD, which were species that were not listed in the WCUPS. Further added was the one species (*Cratylia argentea*) remaining from the Top-830, a species not listed either in the WCUPS or GlobalTreeSearch. Uses for these five species were inferred from the AFD or the World Economic Plants from the USDA GRIN database ^(26)^ (https://npgsweb.arsgrin.gov/gringlobal/taxon/taxonomysearchwep accessed June 2022) and then matched with the use categories of the WCUPS.

Our reason for adding these 67 species was to offer a wider suite of useful woody species to users of GlobUNT, such as widely planted bamboo and rattan species. Users that wish to exclude the additional species and restrict to those included in GlobalTreeSearch can do so in GlobUNT via specific query options. The additional species were also given a different format in the species lists shown in the database. All the additional species have a taxonomic status of ‘ACCEPTED’ in World Flora Online ^(32)^ (WFO), except for the unchecked *Citrus bergamia* uniquely identified as wfo-0000748570 in WFO.

For the additional species not listed in GlobalTreeSearch, the native distribution was obtained from Plants of the World Online (https://powo.science.kew.org; accessed in May and June 2022). For all other 13,951 species, country distributions were obtained by combining the individual country lists from GlobalTreeSearch.

### Species matching

Species matching between species listed in GlobalTreeSearch and the WCUPS was done after preparing standardized lists of species names for the 2022 version of the Agroforestry Species Switchboard ^(23)^. Standardization of nomenclatures for the *Switchboard* was achieved via the *WorldFlora* R package ^(46)^ (version 1.10) by matching names to World Flora Online ^(32)^ (version 2021.12 downloaded from http://www.worldfloraonline.org/downloadData) and, for species that were not matched to WFO, to the World Checklist of Vascular Plants ^(47)^ (WCVP; version 8 downloaded from http://sftp.kew.org/pub/data-repositories/WCVP/).

Matching was thus done separately between GlobalTreeSearch and the *Switchboard* and between the WCUPS and the *Switchboard*, therefore using the master list of standardized names from the *Switchboard* as taxonomic backbone data. This allowed for a straightforward process of including hyperlinks for every species of GlobUNT to the *Switchboard*.

GlobUNT provides information on the type of taxonomic matches for each species, allowing users to verify the credibility of the matches, for example by visually inspecting the similarity in spellings and naming authorities. As hyperlinks are also provided to the matched species names in WFO or WCVP, users can further check for possible changes in taxonomy with the current online versions of WFO and WCVP.

GlobUNT differentiates between six types of taxonomic matches, including (a) direct matches; (b) manual matches; (c) direct matches via WCVP; (d) manual matches via WCVP; (e) direct matches via POWO; and (f) manual matches via POWO.

A ‘direct match’ indicates that the exact name was matched between two species lists. For the 14,014 species listed in GlobUNT, 13, 875 were directly matched with the Switchboard. A ‘manual match’ indicates that a fuzzy match (a match with Levenshtein Distance > 0; see ^(46)^ for details) was accepted after visual inspection. There were 129 of such GlobUNT-Switchboard matches. Matches of ‘direct via WCVP’ or ‘manual via WCVP’ indicate that the species was first matched with a synonym in the WCVP, where the accepted name for the species identified by WCVP was also listed in WFO. There were ten GlobUNT-Switchboard matches of these types, including three manual ones.

For the 14,009 species in GlobUNT where information was obtained from WCUPS, there were 13,856 direct matches, 138 manual matches and one ‘manual via WCVP’ match.

Matches of ‘direct via POWO’ or ‘manual via POWO’ indicate that matching was done via a synonym listed in the Plants of the World Online; 14 of these matches were done, including only one ‘manual via POWO’ match. All ‘matches via POWO’ were done for the WCUPS and for 13 of these, these matches were for the additional species that were not listed in GlobalTreeSearch (additional Table 6; the one exception was *Cupressus lusitanica* matched with *Hesperocyparis lusitanica* in WCUPS).

### Threat status

Information on threat status of individual tree species have been collated through the ongoing Global Tree Assessment ^(18)^ (GTA; https://www.globaltreeassessment.org/) and were obtained from the GlobalTree Portal ^(19)^ (https://www.bgci.org/resources/bgci-databases/globaltree-portal/ accessed on 28^th^ May 2022) by downloading lists for each country.The GTA assigned tree species to the IUCN Red List categories of Extinct, Critically Endangered, Endangered, Vulnerable, Near Threatened, Least Concern, Data Deficient and Not Evaluated. We verified that a unique Red List category had been assigned to species that occur in more than one country.

### Mapping

Countries and territories were allocated to continents and ‘subcontinents’ based on their hierarchical structure within the second edition of the World Geographical Scheme for Recording Plant Distributions

^(48)^ (WGSRPD). This scheme was modified for GlobUNT by:

a. Cape Verde and Saint Helena, Ascension and Tristan da Cunha were assigned to a newly created subcontinental level for Africa of ‘Atlantic Ocean’;
b. Turkey was assigned to Western Asia only (‘Turkey-in-Europe’ was thus ignored); and
c. The Russian Federation was included as a subcontinental level both for Europe and temperate Asia.

Furthermore, the United States Minor Outlying Islands were assigned to different levels within the Pacific based on assignments of individual islands of Johnston I., Midway I., Palmyra I. and Wake I.

Maps were created via the ggplot2 ^(49)^ (version 3.3.6) and sf ^(50)^ (version 1.0-8) R packages using R ^(51)^ (version 4.2.1). Country boundaries were obtained from a Natural Earth ‘admin_0’ vector layer at 1:110 million scale downloaded as a shapefile on 25^th^ September 2022 from https://www.naturalearthdata.com/downloads/110m-cultural-vectors/. The shapefile was processed in QGIS ^(52)^ (version 3.22.11) to split the multipolygon for France into separate polygons for France and French Guyana, and to split the multipolygon for Norway into separate polygons for Norway and Svalbard and Jan Mayen. Further processing was also done to include Kosovo into Serbia, to merge Somalia with Somaliland and to merge polygons for Cyprus. These splits and merges were required to match the country and territory distribution employed by GlobalTreeSearch.

### Calculations of species richness at different levels of geographical aggregation

Species richness at global, continental, subcontinental and country levels were calculated via the dplyr package ^(53)^ (version 1.0.10). Internally in the database, the same package is used to create summary tables of the distribution of species, genus and family richness at different geographical levels.

Data for Brazil and China that have names listed both at the subcontinental and country level in GlobUNT (as in the WGSRPD), were listed once only (for the country level) in tables with the twenty subcontinents and countries with highest richness.

## Data availability

GlobalUsefulNativeTrees can be accessed from https://patspo.shinyapps.io/GlobalUsefulTrees/.

## Acknowledgments

We greatly appreciate funding provided by the Darwin Initiative to project DAREX001 of Developing a Global Biodiversity Standard certification for tree-planting and restoration and by Norway’s International Climate and Forest Initiative through the Royal Norwegian Embassy in Ethiopia to the PATSPO project on Provision of Adequate Tree Seed Portfolio in Ethiopia. We thank Ian Dawson for his comments on earlier drafts of the manuscript and we also thank Kirsty Shaw, Katharine Davies and Galena Woodhouse for their useful suggestions on improving earlier drafts of GlobUNT.

## Supplementary information

### Supplementary Tables

**Supplementary Table 1**. Species richness, genus richness and family richness for GlobalUsefulNativeTrees at country, subcontinental, continental and global levels across the different use categories.

**Supplementary Table 2**. Patterns of endemism for GlobalUsefulNativeTrees at country, subcontinental, continental and global levels.

**Supplementary Table 3**. Patterns of threat status for GlobalUsefulNativeTrees at country, subcontinental, continental and global levels.

**Supplementary Table 4**. Presence of GlobalUsefulNativeTrees species in 27 information sources listed by the Agroforestry Species Switchboard.

**Supplementary Table 5**. Country pledges for the Bonn Challenge (https://www.bonnchallenge.org/pledges) with number of tree species in GlobalNativeUsefulTrees in different use categories

**Supplementary Table 6**. Identities and matching details for the 67 additional species added to GlobalUsefulNativeTrees that were not listed in GlobalTreeSearch

### Supplementary Figures

**Supplementary Fig 1.**
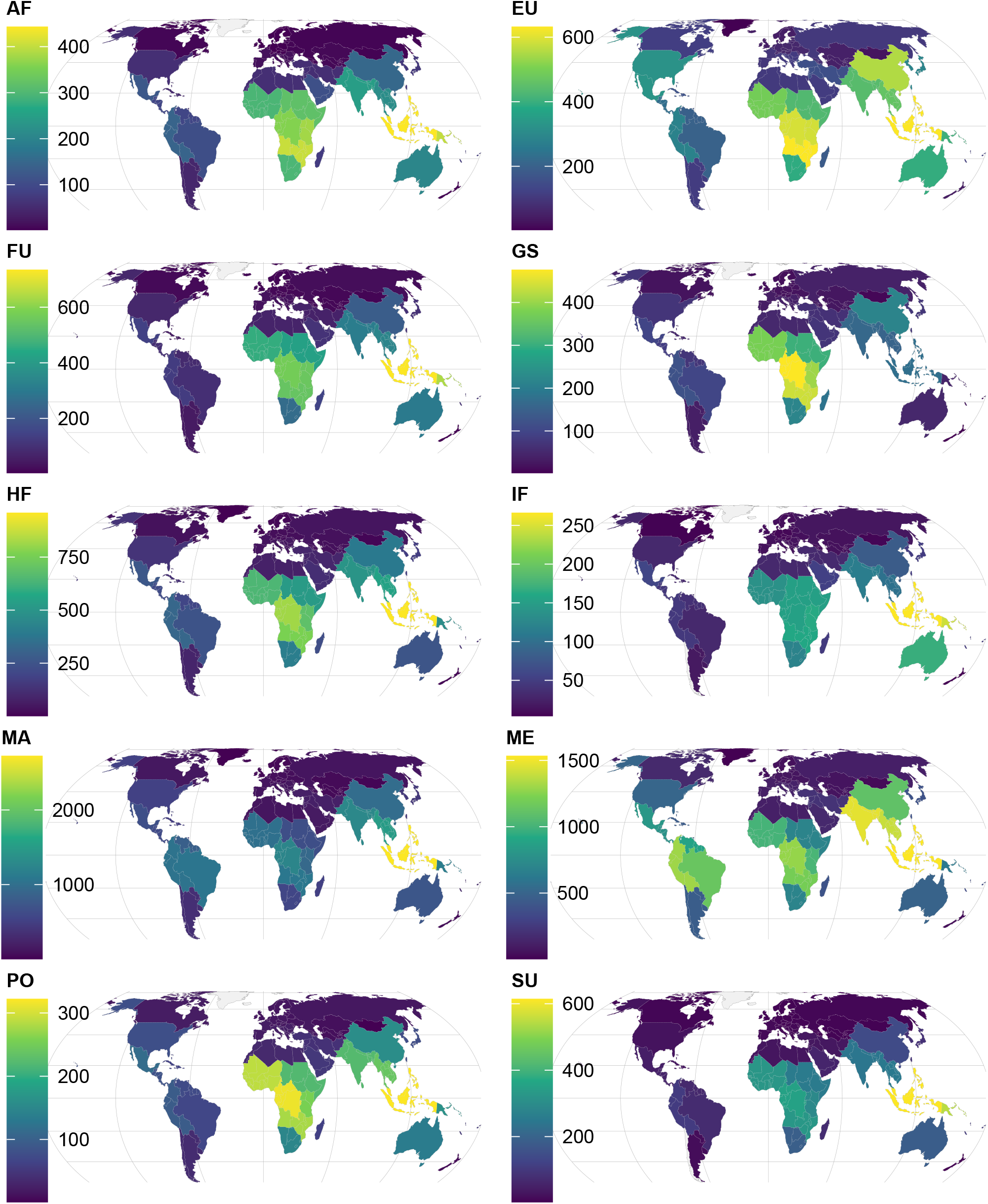
Subcontinental species richness for GlobalUsefulNativeTrees across different use categories. Codes and colour scheme is the same as Fig 1 in the main text, except not to assign the subcontinental colour to a country if that country had no species. Country boundaries added from Natural Earth 1:110 million. Best seen with magnification ≥ 200%.

**Supplementary Fig 2.**
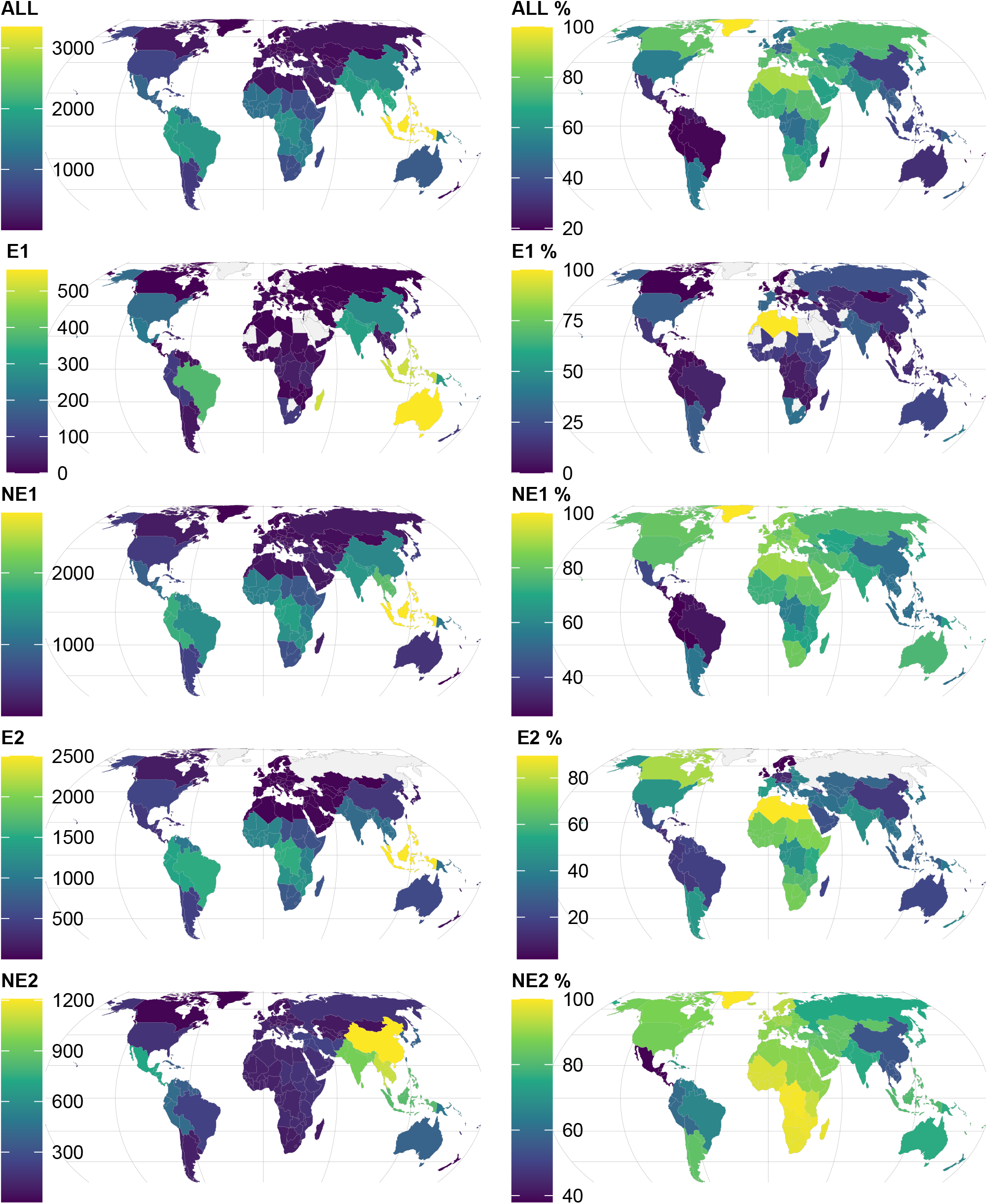
Subcontinental patterns of endemism for GlobalUsefulNativeTrees. Codes and colour scheme is the same as Fig 2 in the main text, except not to assign the subcontinental colour to a country if that country had no species. Country boundaries added from Natural Earth 1:110 million. Best seen with magnification ≥ 200%.

**Supplementary Fig 3.**
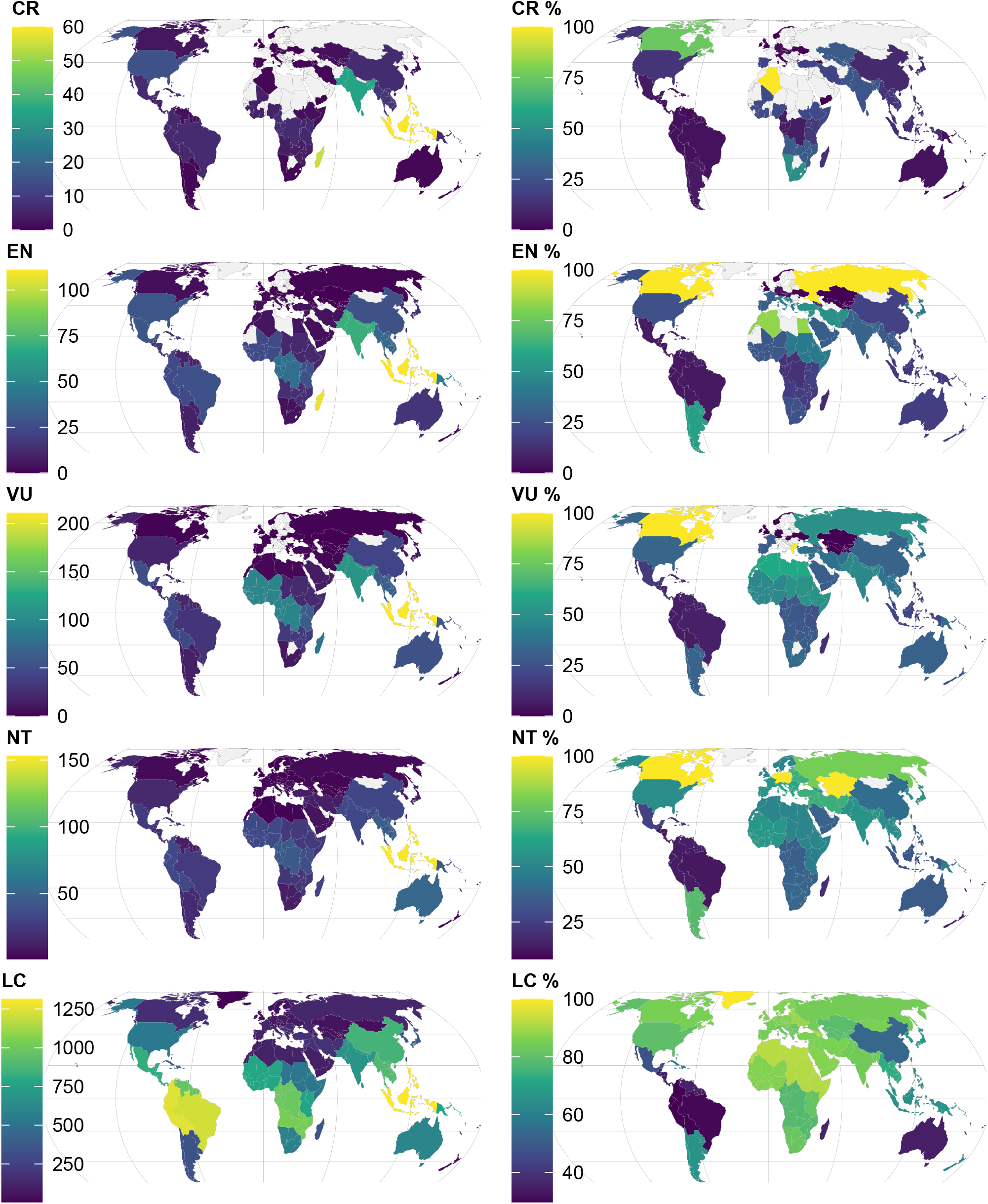
Subcontinental patterns of threats for GlobalUsefulNativeTrees. Codes and colour scheme is the same as Fig 3 in the main text, except not to assign the subcontinental colour scheme to a country if that country had no species. Country boundaries added from Natural Earth 1:110 million. Best seen with magnification ≥ 200%.

